# Simultaneous Super-Resolution and Distortion Correction for Single-shot EPI DWI using Deep Learning

**DOI:** 10.1101/2021.12.03.470880

**Authors:** Xinyu Ye, Peipei Wang, Sisi Li, Jieying Zhang, Yuan Lian, Yajing Zhang, Jie Lu, Hua Guo

## Abstract

Single-shot echo planer imaging (SS-EPI) is widely used for clinical Diffusion-weighted magnetic resonance imaging (DWI) acquisitions. However, due to the limited bandwidth along the phase encoding direction, the obtained images suffer from distortion and blurring, which limits its clinical value for diagnosis. Here we proposed a deep learning-based image-quality-transfer method with a novel loss function with improved network structure to simultaneously increase the resolution and correct distortions for SS-EPI. We proposed a modified network structure based on Generative Adversarial Networks (GAN). A dense net with gradient map guidance and a multi-level fusion block was employed as the generator to suppress the over-smoothing effect. We also proposed a fractional anisotropy (FA) loss to exploit the intrinsic signal relations in DWI. In-vivo brain DWI data were used to test the proposed method. The results showed that the distortion-corrected high-resolution DWI images with restored anatomical details can be obtained from low-resolution SS-EPI images by taking the advantage of high-resolution anatomical images. Additionally, the proposed FA loss can improve the image quality and quantitative accuracy of diffusion metrics by utilizing the intrinsic relations among different diffusion directions.

## I. Introduction

Diffusion weighted imaging (DWI) probes the motion of water molecules and provides valuable information for neuroscience study and diagnosis of diseases [1]. Single-shot echo-planar imaging (SS-EPI), owing to its fast acquisition speed, is widely used for DWI acquisitions [2]. However, due to its limited bandwidth along the phase encoding direction, field inhomogeneities can cause large phase accumulations and lead to severe geometric distortions. Additionally, SS-EPI has a relatively long readout window even with parallel imaging [3], [4], which causes *T*2* blurring artifacts. These artifacts are further aggravated in high-resolution imaging, limiting the achievable resolution of images in clinical practice.

Efforts have been made to improve the image quality of SS-EPI. For distortion correction, a number of methods have been proposed such as the field-mapping method and the top-up method [5], [6]. The field-mapping method uses a multi-echo sequence to measure the B0 field and correct for geometric distortions. In the top-up method, two EPI image sets acquired using two PE gradients with opposite polarities are subsequently used to estimate the displacement maps. A recently proposed method uses echo planar spectroscopic imaging (EPSI) field-mapping followed by intensity correction [7]. However, the efficacy of these post-processing methods relies on the accurate estimation of the displacement maps, especially in regions with severe field inhomogeneities.

A more straightforward way to reduce distortions and improve resolution is to use modified acquisition strategies and shorten the echo spacing (ESP). Multi-shot EPI using interleaved or segmented acquisition trajectories can reduce the effective ESP by a factor of the shot number, thus mitigating the problems for SS-EPI at the cost of scan time [8-10]. Point-spread-function encoded EPI (PSF-EPI) has been proposed to achieve distortion-free high-resolution images by exerting an additional stepwise phase encoding gradient before EPI readout in the conventional 2D SS-EPI [11-14]. To accelerate PSF-EPI, parallel imaging and partial Fourier undersampling can be also integrated. In addition, tilted-CAIPI undersampling and reconstruction strategy can achieve further acceleration (>20 folds) resulting in only 6-8 shots for a 244×244 sampling matrix [15]. However, these multi-shot acquisition strategies are still not efficient enough for clinical high-resolution diffusion tensor imaging or high-angular resolution diffusion imaging [16], [17].

For clinical practice, it is crucial to improve the image quality of SS-EPI while maintaining the acquisition efficiency. Deep-learning-based methods offer a beneficial alternate for this and they have been applied for both distortion correction and high-resolution imaging without sacrificing scan time. For distortion correction, Liao *et al*. [18] designed a convolution neural network (CNN) to correct distortions using a simulated dataset with input distorted images generated from reference distortion-free images. However, it may fail when the field inhomogeneity patterns are different. Hu *et al*. [19] proposed a Unet-based method for SS-EPI distortion correction, where high-quality distortion-free PSF-EPI images were used as references to correct distorted SS-EPI images. This method outperformed the field-mapping and top-up methods. Nevertheless, these deep-learning-based methods cannot improve the image resolution directly and require high-resolution SS-EPI as input, which may prolong the scan time. Meanwhile, deep-learning-based super-resolution method provides a feasible way to generate images with super-resolution from lower resolution input. Dong *et al*. [20] proposed Super-Resolution Convolutional Neural Network (SRCNN) in 2016, through which an end-to-end training process was introduced to learn the mapping functions between the input low-resolution and the target high-resolution patches. Since then, various deep learning-based single-contrast super-resolution methods using different network structures have been explored [21-23]. Several advanced techniques such as channel attention and Generative Adversarial Networks (GAN) have been also employed to improve the performance [24-26]. Recent studies have also demonstrated the feasibility of applying deep learning-based super-resolution methods to boost MRI spatial resolution. Chaudhari *et al*. [27] proposed a 3D CNN network to generate thin-slice knee MR images from thicker input slices. Pham *et al*. [28]and Chen *et al*. [29] implemented 3D CNN network on brain MRI data. Shi *et al*. [30]and Chen *et al*. [31] further proposed improved network structures for MRI super-resolution computation. Other imaging techniques like DWI and dynamic imaging have also been explored [32], [33]. Apart from single contrast-based method, image quality transfer across multiple contrasts has also been introduced by using similar structure information among different MRI contrasts [34], [35].

Based on previous investigations, we hypothesize that deep learning-based methods have the potential to increase resolution and correct distortions simultaneously while maintaining the favorable acquisition efficiency of SS-EPI. Hence, this study aims to generate high-resolution distortion-corrected DWI images from low-resolution SS-EPI images directly through a deep learning network. In particular, we utilized anatomical T2 images as guidance to improve resolution and suppress distortions of DWI images. Additionally, to suppress the smoothing effect from the network, we used a modified GAN method. A dense net with gradient map guidance and a multi-level fusion block is used as a generator to preserve the structure information. Furthermore, we proposed a novel fractional anisotropy (FA) loss to utilize the high-order relations among different diffusion directions. We hypothesize that utilizing such relations can better maintain the image contrasts and improve the quantitative accuracy of measured diffusivities.

## II. Theory

### A. Distortion correction with PSF acquisition

In an EPI acquisition, field inhomogeneity leads to extra phase accumulations in k-space, which induces signal displacements along the phase encoding (PE) direction in the image domain. At a given PE step *k*_*y*_, the corresponding echo time can be calculated by:

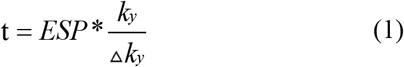

In the PSF-EPI acquisition, an additional phase encoding gradient, namely, PSF encoding, is exerted to the conventional PE direction. Thus, the acquired signal can be formulated as [36-38]:

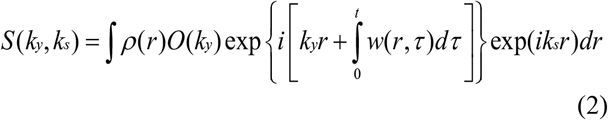

where *k*_*s*_ refers to the PSF encoding, *w*(*r,τ*) is off-resonance which varies with spatial location *r* and time *τ*. *O*(*k*_*y*_) refers to relaxation terms and *ρ*(*r*) refers to proton density

The inverse Fourier transform of (2) leads to

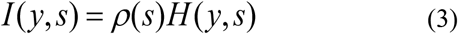

Where

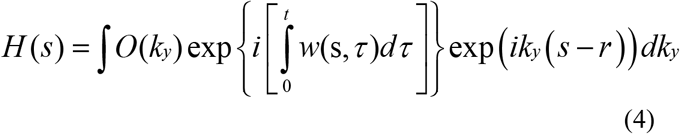

H(s) can be regarded as a PSF function for the acquisition, indicating that how the image is shifted and blurred along the PE direction due to the field inhomogeneities and *T*2 * relaxation.

By an integration along y and s respectively, we can get:

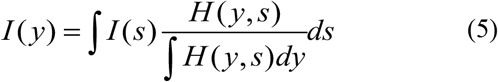

Therefore, the distorted image *I*(*y*) can be described as the convolution of the undistorted image *I*(*s*) with the convolution kernel 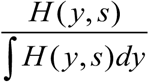 along the PSF encoding direction. Then, undistorted image *I*(*s*) can be obtained by deconvolution from *I*(*y*) [36-40]. Thus, a CNN network can be used to correct the distortions by taking the advantage of multiple convolution layers.

### B. Deep learning-based SR problem

Given a low-resolution image patch *I*_*LR*_ and its corresponding high-resolution image patch *I*_*HR*_, the learning-based super-resolution methods try to find the following mapping function f so that:

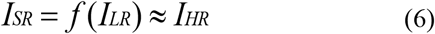

And the ith point of the super-resolution image *I*_*SR*_ can be described as [41]:

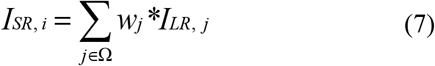

where Ω refers to the corresponding neighborhood of the low-resolution input point. Thus, the reconstruction of the super-resolution image can be viewed as a convolution process of low-resolution image patches.

Recently, deep learning methods like CNN networks have been widely explored for SR reconstruction, since they can better extract the edge information and learn the nonlinear mapping functions between low-resolution and high-resolution patches than the traditional learning-based SR methods which require prior information. Assuming another reference image *I*_*ref*_ is provided, the deep learning method typically tries to minimize the following least-square cost function:

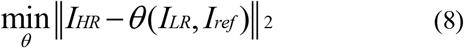

where *θ* refers to the network

Since 2D CNN network can tackle both super-resolution and distortion correction problems, we used a 2D CNN-based network for simultaneous super-resolution and distortion correction to improve the image quality of low-resolution SS-EPI.

## III. Methods

### A. Data acquisition

14 healthy volunteers were recruited and scanned on a Philips Ingenia CX 3T scanner (Philips Healthcare, Best, The Netherlands). All human studies were performed under the institutional review board approval from our institution, and written consents were obtained before the study. Different contrast brain MRI images were acquired with detailed scan parameters shown in Table I.

**TABLE I.** Acquisition protocols.

|  | TR (ms) | TE (ms) | Resolution (mm<br>× mm × mm) | R<sub>PE</sub>/R<sub>Ps</sub><br>F | bvalue<br>(ss·mm<sup>-2</sup>) | Directions | NEX | Time(min:sec) |
| --- | --- | --- | --- | --- | --- | --- | --- | --- |
| PSF-EPI | 6293 | 67 | 1 × 1 × 4 | 4/14 | 0,1000 | 6 | 1 | 08:54 |
| SS-EPI | 3050 | 67 | 1 × 1 × 4 | 3 | 0,1000 | 6 | 4 | 01:32 |
| SS-EPI-opp | 3050 | 67 | 1 × 1 × 4 | 3 | 0,1000 | 6 | 4 | 01:32 |
| ME FFE | 5.5 | 1.03 | 3.44 × 3.44 × 4 | 3 | - | - | 1 | 00:03 |
| T2W-TSE | 3000 | 80 | 1 × 1 × 4 | No | - | - | 1 | 01:24 |
| LR SS-EPI | 3050 | 67 | 2 × 2 × 4 | 3 | 0,1000 | 6 | 2 | 00:46 |

The datasets were sorted into two groups, of which the first one was used for synthesizing training data and the second used for testing. Specifically, the first dataset included 12 subjects. Both SS-EPI and PSF-EPI DWI were acquired with 1mm×1mm in-plane resolution. SS-EPI was acquired with 4 averages to provide 4 training pairs for each subject. PSF-EPI DWI was acquired with 4-fold acceleration along the PE direction and 14-fold acceleration along the PSF encoding direction., which is equivalent to 12 shots for data sampling. The simulated low-resolution SS-EPI input data were obtained by undersampling the high-resolution SS-EPI data to 2mm×2 mm in-plane resolution. The images were first Fourier transformed and then cropped, after which an inverse Fourier transform produced the low-resolution images. To compare with the traditional distortion correction methods, a 3D multi-echo gradient echo sequence was used to obtain field maps and an additional SS-EPI with opposite PE direction (SS-EPI-opp) was also acquired for the top-up distortion correction. 2D T2W-TSE images were acquired to provide an anatomical reference. All sequences shared the same FOV covering the whole brain. The anatomical images and DWI images were registered to reduce mismatch due to motion. 9 subjects in this group were used for training, 2 for validation and 1 for test. The second group included 2 subjects for test. SS-EPI DWI used in-plane resolution of 2mm×2mm whereas PSF-EPI used 1mm×1 mm. The other DWI acquisition parameters were kept consistent in the 2nd group. To be brief, the test dataset included 1 subject with simulated low-resolution SS-EPI and 2 subjects scanned with in vivo low-resolution SS-EPI.

### B. Network structure

In this study, we propose a network to obtain high-resolution distortion-corrected images from low-resolution distorted images directly. So, it attempts to learn the mapping functions between the low-resolution input patches and the high-resolution output patches. Since the distortion occurs along the PE direction mainly, patches are extracted via cutting images along the frequency encoding (FE) direction. The patch size is 112×16 (PE×FE) for the input low-resolution SS-EPI, and 224×32 for the output high-resolution PSF-EPI.

The specific network structure is shown in Fig. 1. The generator consists of four branches, namely feature extraction, gradient estimation, reconstruction and fractional anisotropy (FA) calculation. The feature extraction branch uses a densely connected 2D CNN network with a multi-level fusion block. The input SS-EPI patches are concatenated with the T2W patches. The dense network consists of several residual blocks, and a skip connection exists between each pair of the blocks. Each residual block is composed of three convolutional (Conv) layers followed by ReLU activation and BN layer. Four residual blocks are used in the feature extraction branch. Then all information from different blocks is fused together followed by the reconstruction through a bottleneck structure [42] to save memory burden.

**Fig. 1.**
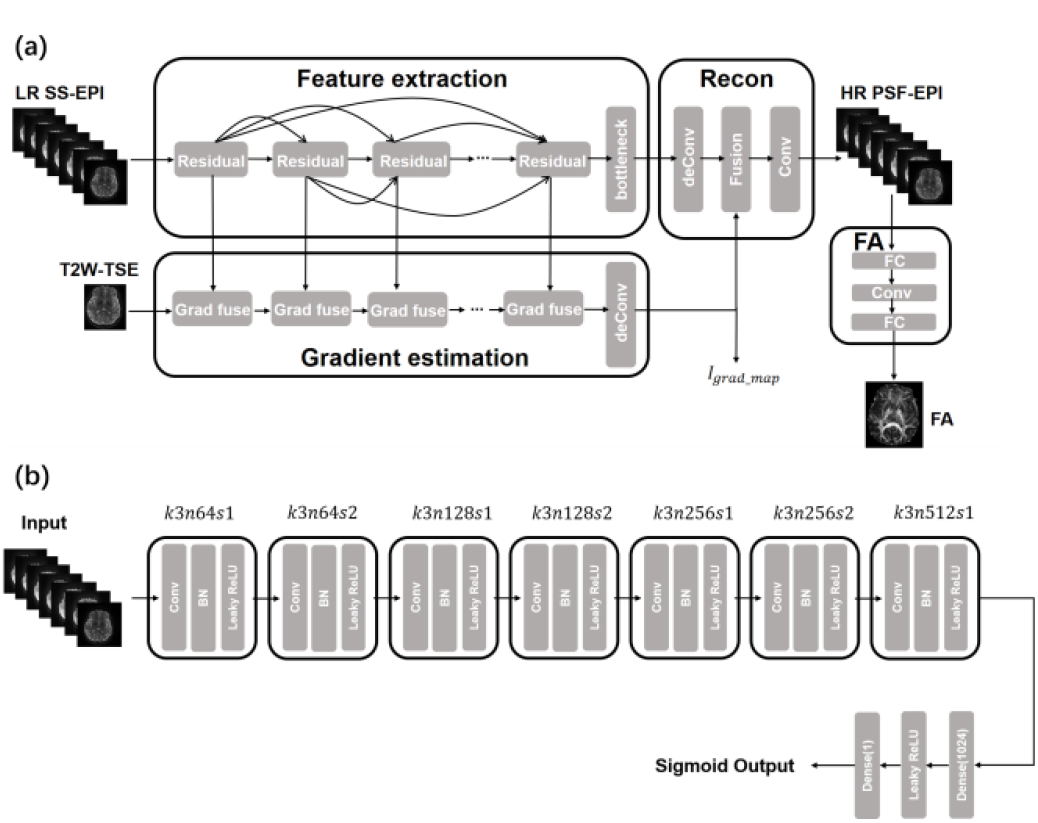
Network structure. (a) The generator has four branches, namely Feature extraction, Gradient estimation, Reconstruction and FA calculation. (b) The discriminator is adopted from SRGAN.

To better preserve the edge and texture information, we use a gradient estimation branch to estimate the gradient maps and guide the image reconstruction [43]. The gradient branch has 4 gradient blocks composed of three Conv layers. A skip connection exists between the initial gradient map estimation and the final output estimation. Each gradient block uses Sobel operator to extract gradient information from feature maps in the feature extraction branch and uses Conv layers to update the gradient map estimation. The final output gradient maps are concatenated with the output of feature extraction branch. Additionally, the output of the gradient branch is also compared to the ground truth gradient maps and serves as gradient map loss.

The reconstruction branch concatenates the estimated gradient maps with multi-level information fused in the bottleneck structure. The last convolutional layer applies seven filters with size 1×1 to obtain the final output images.

The FA calculation branch is designed to introduce our proposed FA loss which will be explained in the following section.

The discriminator is adopted from the previously reported SRGAN [25]. Specifically, the discriminator uses the vgg-19 network structure [44], including 8 conv layers with LeakyRelu activation function. The full connection layer and sigmoid activation function at the end of the network give the probability of the predicted images from real high-resolution images and generated high-resolution images.

Since different diffusion directions share similar structure information, they are jointly used as the network input. Thus, the input images have 8 channels (including 1 b = 0 s/mm^2^ and 6 b = 1000 s/mm^2^ images along 6 diffusion directions from SS-EPI as well as 1 T2W-TSE image) and the output images consist of 7 channels (including 1 b = 0 s/mm^2^ and 6 b = 1000 s/mm^2^ images along 6 diffusion directions acquired by PSF-EPI).

### C. FA loss

In diffusion tensor imaging, FA values, representing the extent of anisotropy of the diffusion ellipsoid, provide microstructure information such as fiber orientation and explore high-order relations between different diffusion directions [45], [46]. Thus, we utilize FA values as an extra loss to improve the quantitative accuracy of diffusion metrics.

FA calculation needs tensor fitting and eigenvalue decomposition which hinders back propagation when directly implemented in the network. Thus, we pre-train a FA calculation network F to calculate FA values from the input DWI images. The network F consists of two FC layers and one CNN layer and is trained on the high-resolution PSF-EPI images. The target FA values are calculated using FMRIB Software Library (FSL) [47]. After pre-training the network F, we fix the weights of F and use it as the FA calculation branch. During the training of the main network, the FA calculation branch calculates FA maps using the super-resolution outputs and the high-resolution targets, respectively. The differences of the FA results are termed as the FA loss:

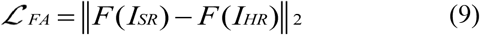

For the loss function design, we combine the structural consistency index (SSIM) [48], pixel-wise difference (ℒ_1_), image edge preservation (Gradient loss and gradient map loss) [49], the proposed FA loss and the cross-entropy GAN loss (ℒ_*GAN*_). The loss function can be written as

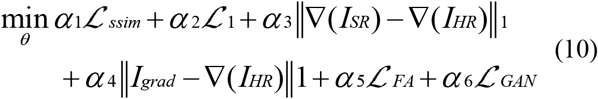

Here ∇ refers to gradient operator, and ℒ_*GAN*_ refers to:

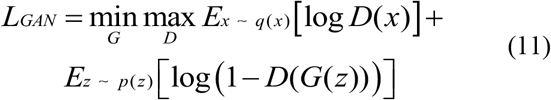

### D. Implementation

At the training and validation stages, the grayscale of T2W-TSE, SS-EPI and PSF-EPI images were normalized to (0,1). Specifically, the DWI images from SS-EPI and PSF-EPI were normalized using the maximum value of b = 0 s/mm^2^ images so that the computation of diffusion metrics would not be affected.

The proposed network was implemented using Keras. Adam Optimizer was used for minimization. The total trainable number was 2.5 million. The batch size was set to 8, and the maximum number of epochs was set to 100. We used one NVIDIA GTX 2080ti GPU for training and validation. The total training time was about 24 hours.

### E. Evaluation

To assess the performance of the proposed network, the results were also compared to those from the traditional distortion correction methods and other deep-learning networks.

First, the proposed method was compared to the field-mapping and top-up methods as well as the previously reported U-net based distortion correction (termed as Unet DC) [19] method. The purpose is to investigate whether the image quality of our method using low-resolution input is comparable to those using high-resolution input.

Then, we compared our method to Bicubic interpolation (LR), Resnet, Unet and SRGAN [25] for joint super-resolution and distortion correction using simulated test data and in vivo test data. Resnet and Unet were trained with a combination of L1 loss, SSIM loss and Gradient loss. The weightings are set the same as in [19]. SRGAN was trained with L1 loss, SSIM loss, Gradient loss and Adversarial loss. The networks trained without GAN and FA loss were also compared. To evaluate the quantitative accuracy, mean diffusivity (MD) and FA values were calculated using FSL. Root Mean Squared Error (RMSE) were used to illustrate the accuracy.

Furthermore, two experiments were conducted to investigate the generalization feasibility of the proposed method. The first experiment was designed to see if the network can deal with different distortion patterns. SS-EPI images with an opposite PE direction (SS-EPI-opp) from another in vivo test case was used as input. Secondly, low-resolution distortion-free images obtained by undersampling PSF-EPI were also used as input to investigate the performance of the proposed network when no distortion was present.

## IV. Results

Fig. 2 shows the results from the pre-trained FA calculation network. The FA maps calculated by the network are compared to the reference FA maps obtained by FSL. Compared to the reference, the results show high consistency. Thus, we can use the FA calculation network to calculate FA values directly from input DWI images.

**Fig. 2.**
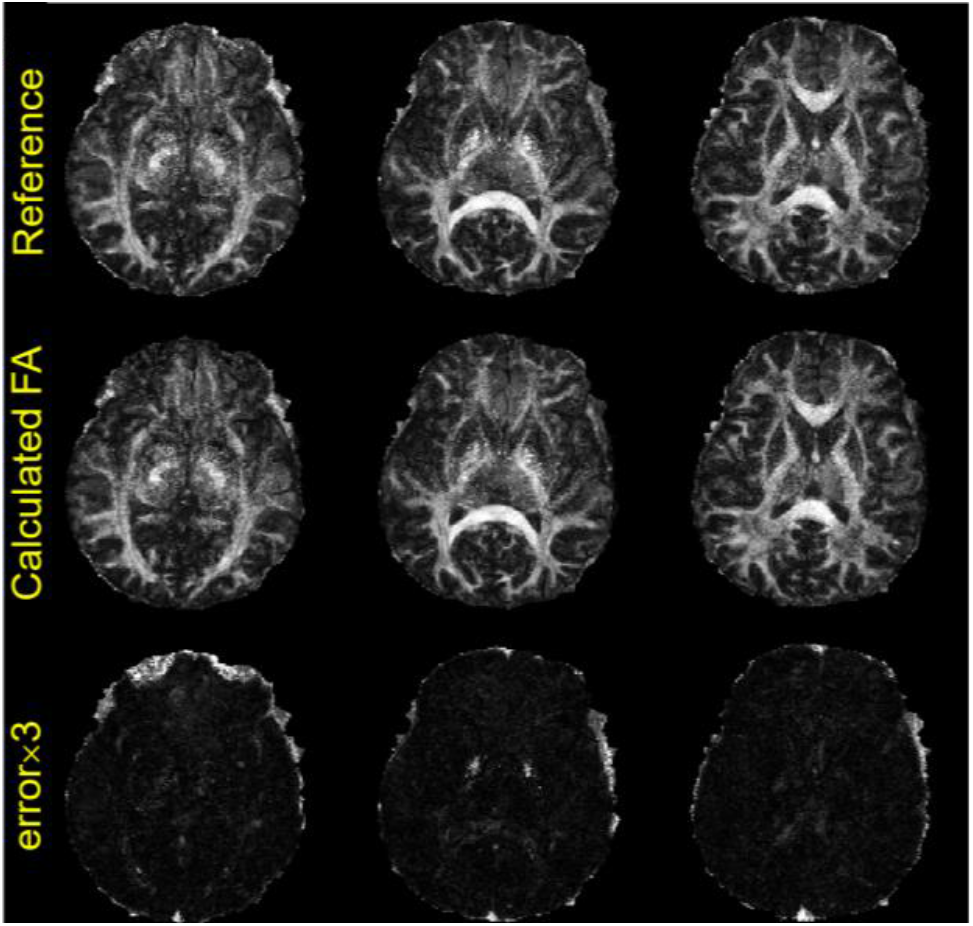
Results of the pre-trained FA calculation network. The first row and the second row show the reference FA maps and the results of the FA calculation network respectively. Error maps are shown. Based on the error maps, the proposed calculation network can calculate the FA values precisely.

Fig. 3 shows the b = 0 s/mm^2^ and mean DWI results for the validation case. Two representative slices of the low-solution SS-EPI, high-resolution PSF-EPI, T2W-TSE and the super-resolution results are shown. Compared to the low-solution SS-EPI, the proposed network can correct for distortions in the regions affected by field inhomogeneities such as the basal temporal regions and the eyeballs. The corrected images in the fourth column show high structural similarity with the reference images. Additionally, as shown in the zoomed-in regions, the low-solution SS-EPI images lose structure details which can be recovered after processing. Meanwhile, the proposed method can recover the sharp edges.

**Fig. 3.**
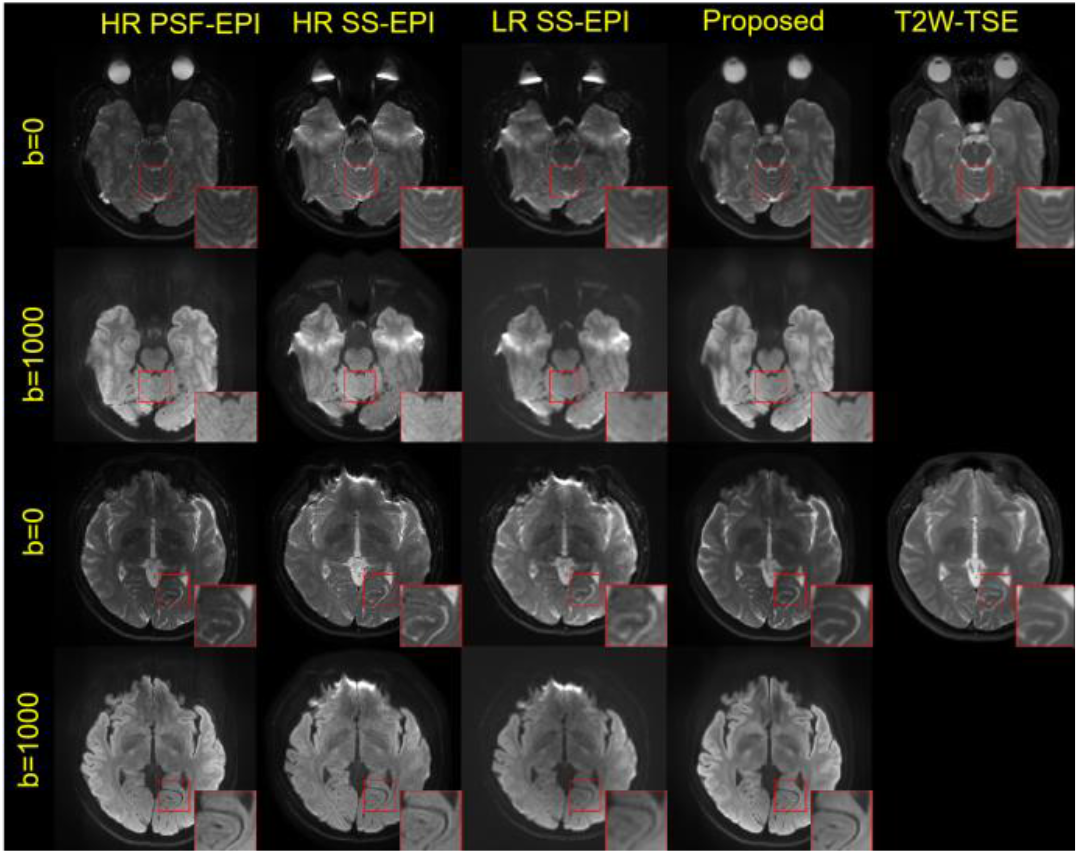
Results of the validation test. Two representative slices of the distortion-free PSF-EPI reference, high-resolution distorted SS-EPI, input low resolution distorted SS-EPI and outputs of the proposed method are shown. Detailed structures are magnified and shown in the corner for better evaluation. Based on the b0 and mean DWI results, the proposed method can correct the distortions presented in the SS-EPI images and increase the resolution, recovering detailed structures compared with the low-resolution input. The recovered structures are consistent with T2 images.

To demonstrate the potential to replace high-resolution acquisition with low-resolution acquisition, Fig. 4 compares the proposed method with other distortion correction methods. The proposed method uses low-resolution images as the input and the others use high-resolution images. High signal intensities still exist due to the estimation errors of the displacement maps after field-mapping and top-up correction. Moreover, the results are blurry in the regions pointed by the red arrows thus the edge details are hardly seen, whereas the proposed method successfully resolves the signal pileups and the recovered structures show higher consistency with the distortion-free references. Although using low-resolution images as the input, the proposed method shows better image quality than the field-mapping and top-up methods in the regions pointed by the red arrows. Compared to Unet DC using high-resolution SS-EPI as the input, the proposed method using low-resolution SS-EPI still demonstrates comparable perceptual quality.

**Fig. 4.**
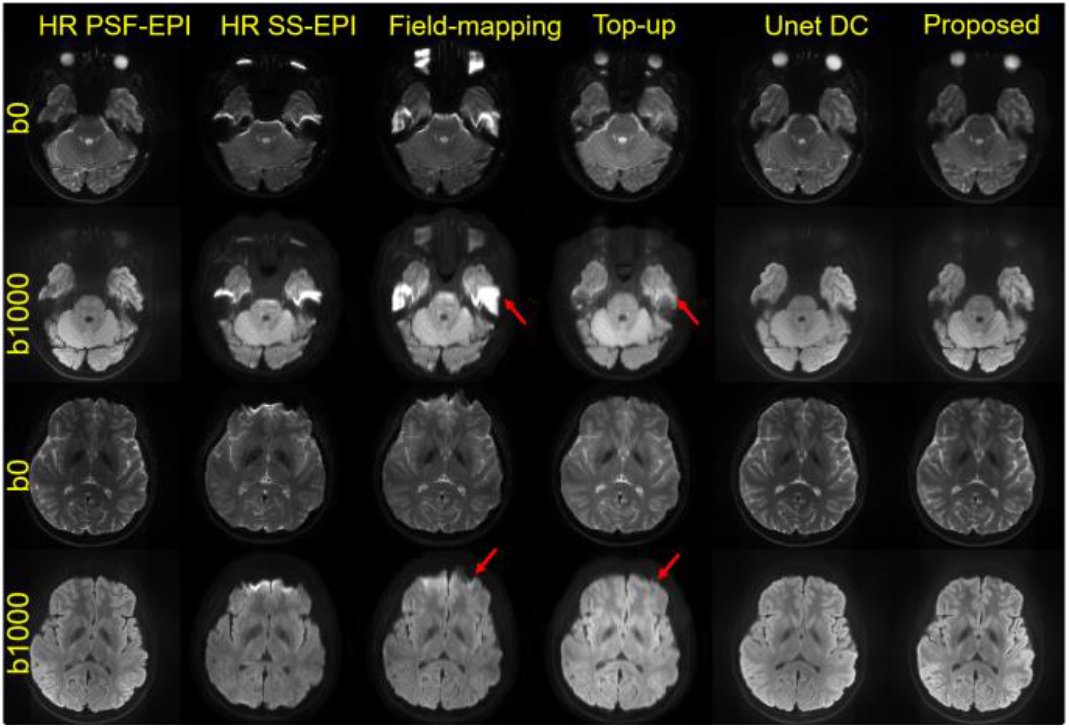
Comparison between the proposed method and the traditional distortion correction methods for high-resolution images along with the Unet-DC method using high-resolution input for the simulated test case. Two representative slices of b0 and mean DWI images are shown. The red arrows indicate the regions where remaining distortion artifacts exist. The proposed method outperforms the traditional distortion correction methods and has comparable image quality with the Unet-DC method using high-resolution input, indicating that we can directly use low-resolution input images to recover distortion-free high-resolution images.

The comparison between the proposed network and other deep-learning networks using low-resolution distorted input images is shown in Fig. 5. Mean DWI images are shown. As pointed by the arrows, the Unet method shows smoothed texture information. Although the distortion can be corrected, the details are hardly seen. Resnet and SRGAN both show better sharpness compared to Unet, but still suffers from inaccurate edge information in the recovered regions. The w/o FA and w/o GAN methods show visually more similar images in the recovered regions compared to previous networks whereas the proposed method preserves more detailed features compared with the distortion-free references. This comparison shows the gradient map guidance and FA loss can help improve the image quality.

**Fig. 5.**
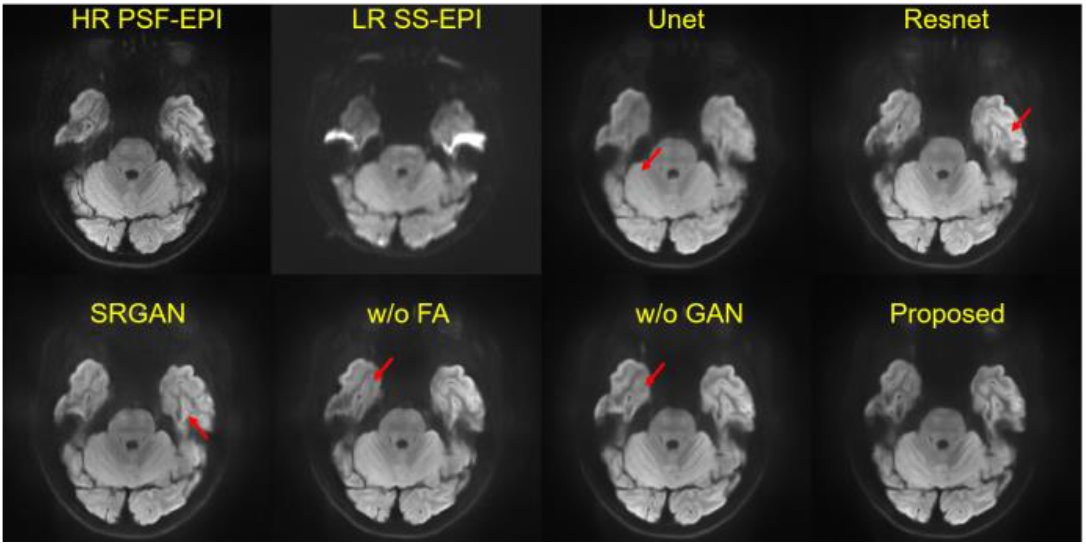
Comparison between the proposed method and other deep learning methods for the simulated test case. Mean DWI results of different networks for simultaneous super resolution and distortion correction for low-resolution SS-EPI are shown. Red arrows refer to the regions which are not well recovered. Compared to the proposed method, other methods lose detailed structure information as pointed by the red arrows.

The efficacy of distortion correction is also evaluated in the color-coded FA maps. As shown in Fig. 6, compared to the SS-EPI, the proposed method improves the overall image quality. The color patterns are consistent and the proposed FA loss can improve the white matter fiber pathway as pointed by the red arrow. Meanwhile, the proposed network without FA loss shows better preserved grey matter structures.

**Fig. 6.**
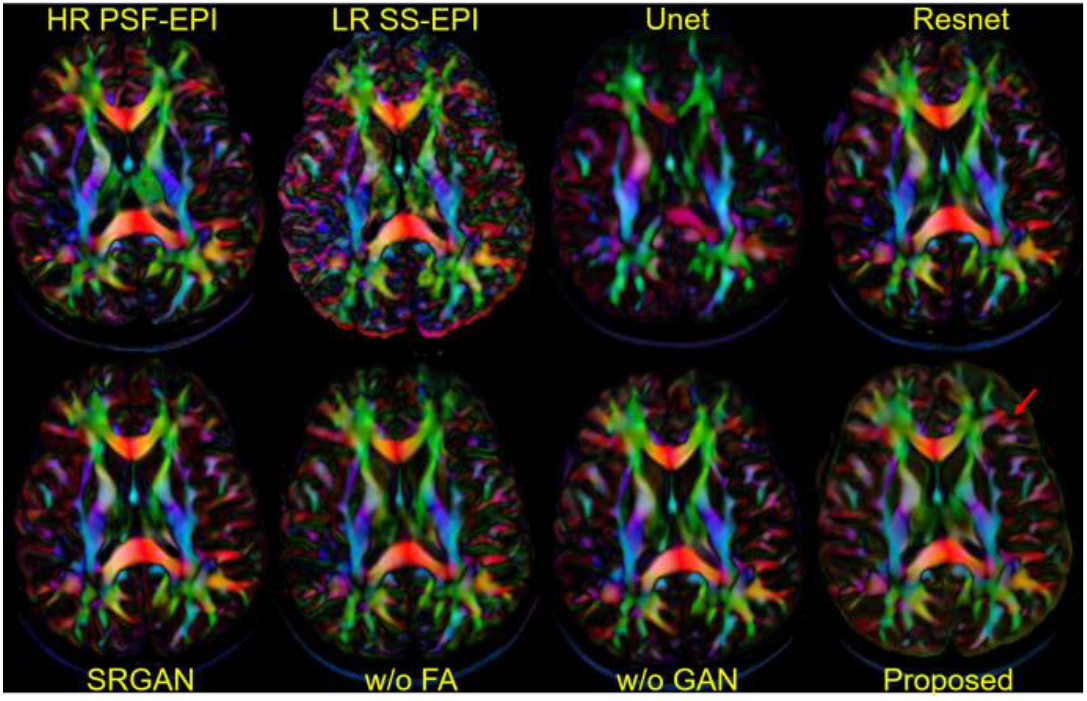
Color-coded FA maps obtained from different simultaneous super resolution and distortion correction methods. Compared to the referenced cFA maps, only the proposed method can recover the white matter fibers as pointed by the red arrow.

The errors of measured diffusion metrics between the reconstructed images and PSF-EPI images are compared in Fig.7. Quantitative results of whole-brain MD and FA errors are shown in Fig. 7 (a) and Fig. 7 (b) respectively. Compared to the other deep learning methods, as expected, the proposed method trained with FA loss showed better accuracy, further indicating that the proposed FA loss can better improve the quantitative accuracy by maintaining the anisotropic relations among different diffusion directions.

**Fig. 7.**
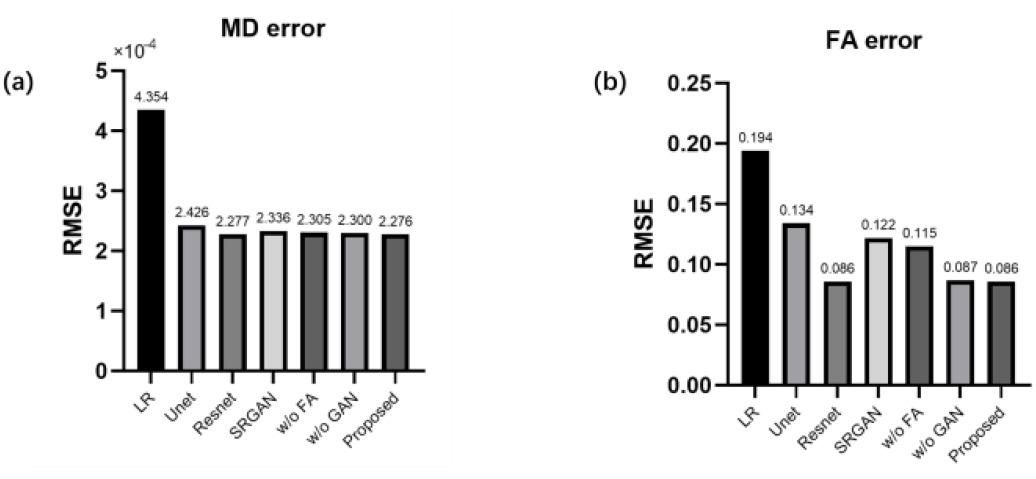
Comparison of quantitative DTI metrics obtained from different simultaneous super resolution and distortion correction methods using High-resolution distortion-free PSF images as references. (a) Whole-brain RMSEs for MD, and (b) Whole-brain RMSEs for FA.

Fig. 8 shows the results using in-vivo low-resolution test data as the input. The high-signal intensities caused by signal piling-up are corrected by the proposed method. In the enlarged views of the red boxes, fine details and preserved edges consistent to the reference images can be observed. Compared to the network trained without GAN or FA loss, the proposed method can suppress the over-smoothing effects. The textures and the curves become much clearer and detectable in general, illustrating the improvement of the perceived resolution.

**Fig. 8.**
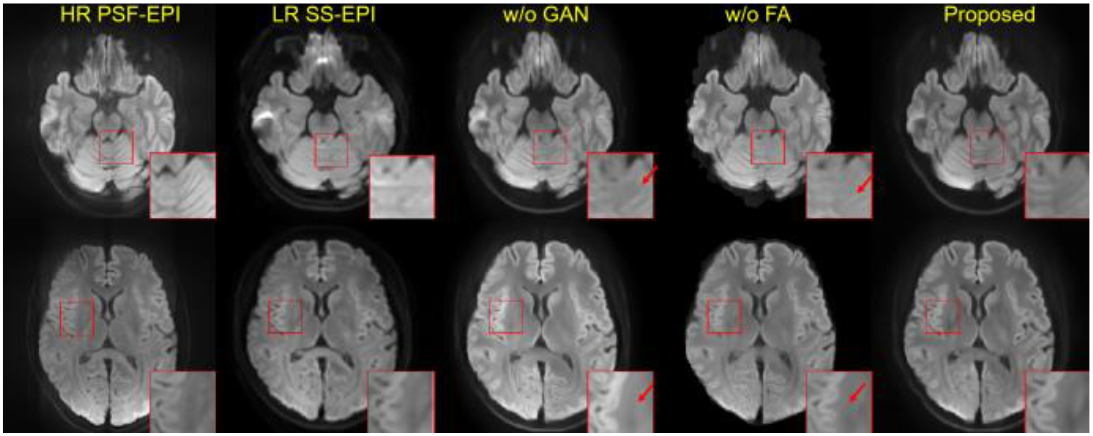
Joint distortion correction and super resolution reconstruction results of the in vivo low-resolution SS-EPI. Mean DWI images are provided. The enlarged view of tissue boundaries is shown. Red arrows refer to the areas where detailed structure information is lost. Based on the results, the proposed method demonstrates the best consistency with the reference images.

Since the proposed method used the same PE direction during training, we further tested the data from another in vivo test case acquired with opposite directions (SS-EPI-opp). As shown in Fig. 9, though a few residual artifacts exist, the proposed method can still correct major distortions such as the stretched eyeballs and improve the perceived resolution for the opposite direction. The proposed method also better preserved the fine structures as pointed by the red arrows.

**Fig. 9.**
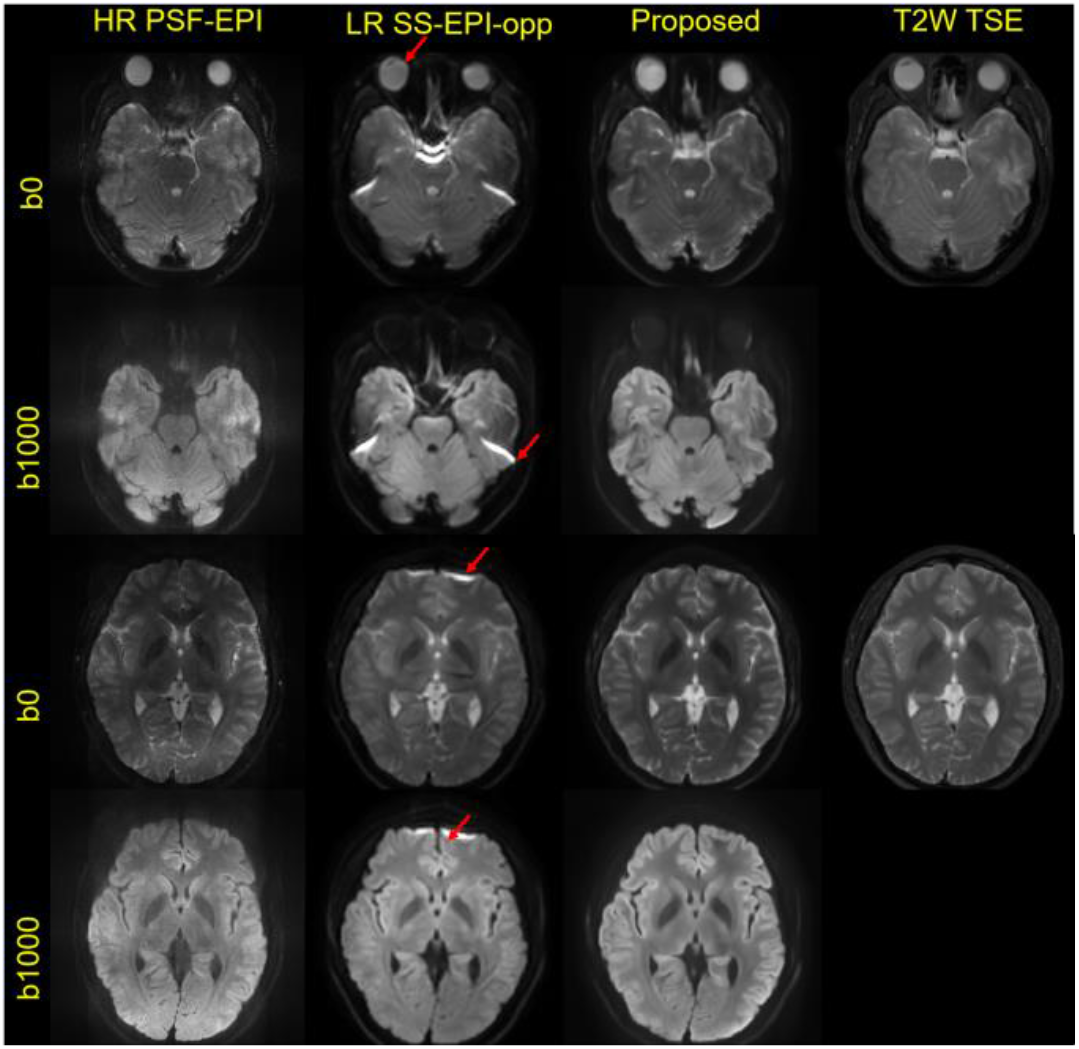
Results for SS-EPI acquired with opposite directions of another in vivo test case. The b0 and mean DWI and outputs of the proposed method are shown. Red arrows refer to the regions where the distortion patterns are different. Though the distortion patterns are different, the proposed method still manages to correct the distortions and increase the resolution, providing more detailed information.

To further illustrate that our proposed method can recognize the distortion patterns, as shown in Fig. 10, we undersampled the PSF-EPI data by 2×2 in-plane factor as undistorted low-resolution input, and found that the network managed to clearly recover the detailed tissue boundaries whereas the curves were difficult to be seen in the low-resolution image. At the same time, the distortion-free geometric patterns were maintained.

**Fig. 10.**
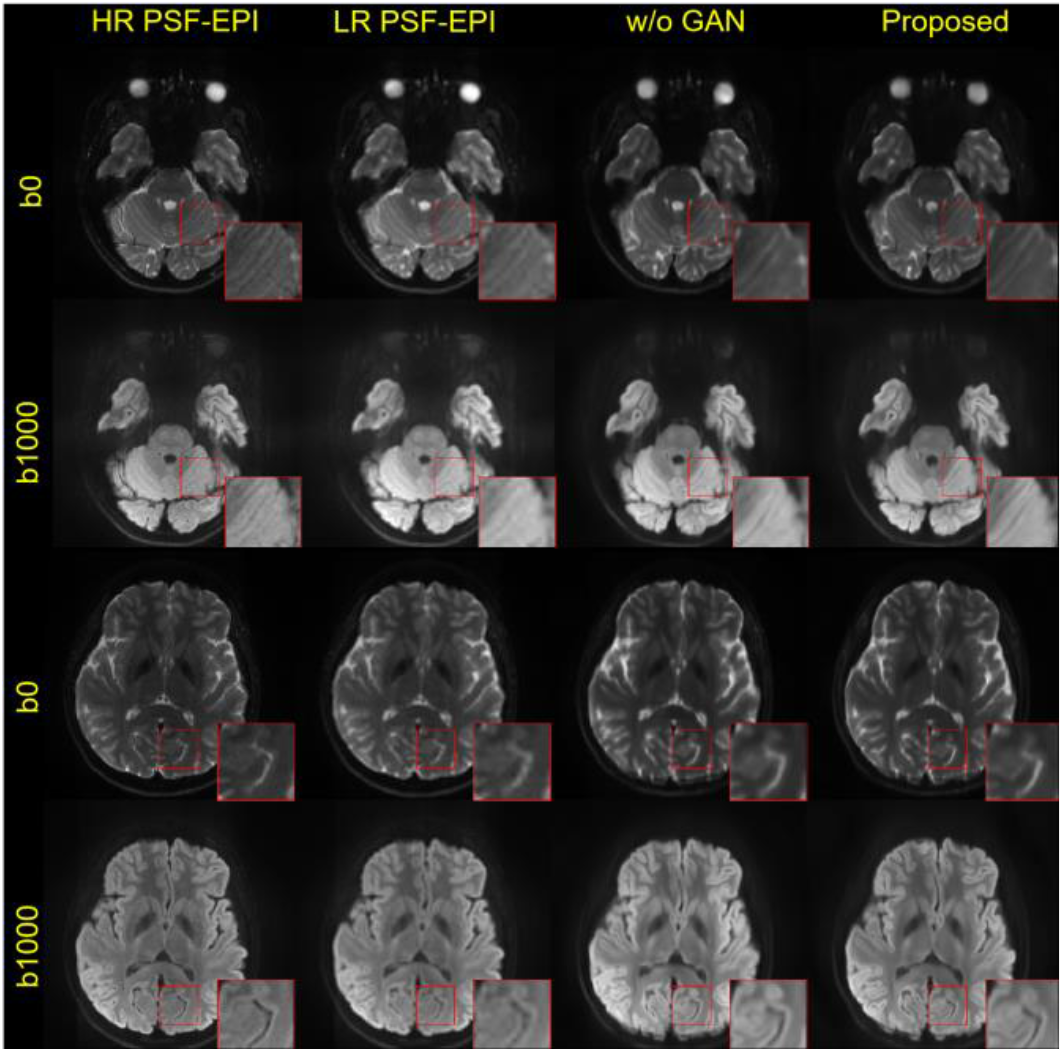
Results of the simulated low-resolution distortion-free PSF-EPI images. The b0 and mean DWI of reconstructed super-resolution images from the proposed method are shown. Detailed structures are magnified and shown in the corner for better evaluation. When no distortion, the proposed method keeps the original geometric information while increases the resolution.

## V. Discussion

SS-EPI is widely used for clinical DWI acquisition, yet even if parallel imaging is employed, the limited bandwidth along PE direction still leads to severe distortions and blurring artifacts for high-resolution acquisition. Advanced EPI-based techniques including multi-shot EPI and PSF-EPI techniques can mitigate these problems and improve the image quality. However, with higher resolution, the prolonged scan time renders these techniques impractical for clinical examinations, especially for emergencies. Post-processing methods can improve the image quality while maintaining the sampling efficiency. However, conventional post-processing methods heavily rely on the accurate estimation of the displacement maps which might be difficult to achieve given limited scan time.

In this study, we propose a deep-learning-based method to simultaneously correct distortions and increase perceived resolution for low-resolution SS-EPI. Compared to previous works, this network only requires low-resolution input to recover high-resolution distortion-free diffusion images. The preliminary results show that the proposed method outperforms the state-of-the-art distortion correction methods using high-resolution input. Additionally, compared with other deep learning methods, the proposed method further increases the reconstruction accuracy of the detailed structures with the gradient map guidance. Quantitative results also show high consistency between the proposed method and the ground truth. The results indicate that the proposed FA loss improves the accuracy of diffusion metrics by maintaining the contrast variations between different diffusion directions.

Diffusion contrast plays a vital role to obtain accurate quantitative measures. Different diffusion directions share the same structure information yet the contrasts vary. The pixel-wise based loss might lead to smoothing. Based on the results, the proposed method with FA loss can better recover the detailed anatomical structures and the calculated diffusion modeling parameters are more accurate, indicating that the FA loss can capture the intensity variations between different directions. However, though the improvement of white matter fibers in color-coded FA maps is obvious with FA loss, the grey matters are suppressed compared to the results without FA loss. One potential reason is that the FA values of white matter are higher than that of grey matter, so the FA loss attaches more importance to the former. FA loss with decayed weight during training might be helpful to recover both white matter and grey matter. which will be our further work. Though the FA loss needs fixed 6-direction diffusion images as input, we can duplicate the directions to fill in 6 channels when the number of directions is not enough.

Since we use 2D acquisition protocols to cover the whole brain with 25 slices, a 2D network is used instead of commonly used 3D net to achieve super-resolution. Besides, this work demonstrates the potential of the proposed network to simultaneously increase perceived resolution and correct distortions, more comprehensive comparison with other advanced network structures is beyond the scope of this paper. Despite the performance of the proposed method, there are still several limitations. Firstly, our network used a relatively small number of training samples without patients’ data. Additionally, the training data was acquired with the same PE direction under the same protocol for brain. Data for fine-tuning is needed to get more accurate results for different protocols or different b values. Thus, to further increase the generalizability, patient data with different diseases and lesion types acquired using different protocols can be included in the training process. Secondly, the FA loss was trained on fixed diffusion directions and extension to other patterns of diffusion directions needs retraining. Further study can use quantization methods to extend the network to different diffusion directions settings. Finally, the proposed method shows smoother FA results and the mean DWI images still show slight variations with the ground truth, since the network only uses low-resolution input and CNN can lead to smoothed results. The adaptation of loss functions in the future can help produce sharper and more reliable quantitative results. In the future, we will investigate other methods like including loss functions for diffusion tensor values or adding extra anatomical references like FLAIR to improve the performance.

## VI. Conclusion

In conclusion, we propose a deep learning-based method to simultaneously increase the perceived resolution and correct distortions for low-resolution SS-EPI, thus the high-resolution distortion-free images can be obtained directly from low-resolution distorted images with the guidance of anatomical images. The results demonstrate that the proposed method outperforms the traditional distortion correction methods and shows improved image quality compared with other deep learning networks. Furthermore, the proposed network also demonstrates the potential to obtain high-quality DWI images for lesion detection in clinical routine scan with practical acquisition time.

## Appendix

See Table II

**TABLE II.**
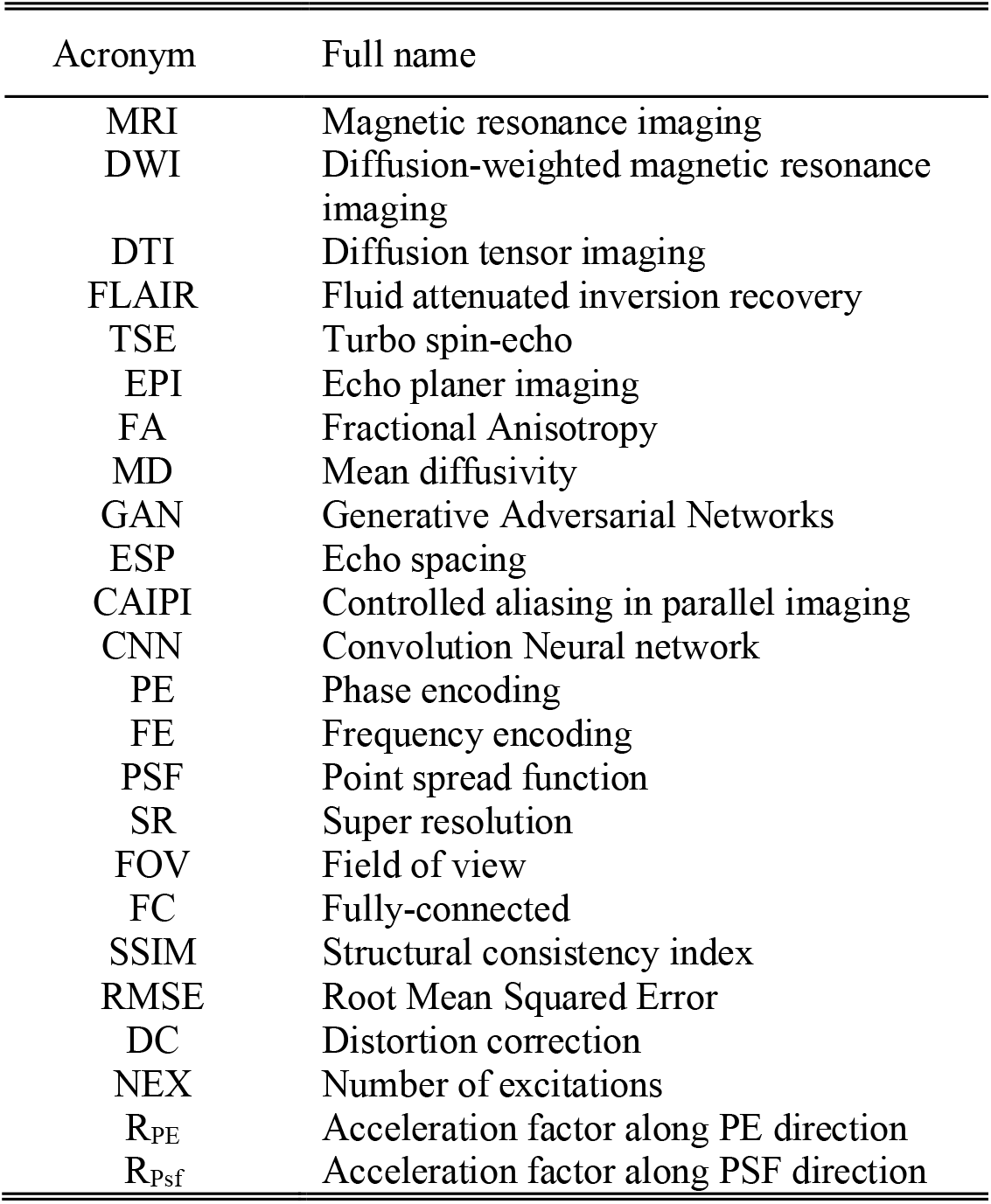
List of all the Acronym in this paper

| Acronym | Full name |
| --- | --- |
| MRI | Magnetic resonance imaging |
| DWI | Diffusion-weighted magnetic resonance imaging |
| DTI | Diffusion tensor imaging |
| FLAIR | Fluid attenuated inversion recovery |
| TSE | Turbo spin-echo |
| EPI | Echo planer imaging |
| FA | Fractional Anisotropy |
| MD | Mean diffusivity |
| GAN | Generative Adversarial Networks |
| ESP | Echo spacing |
| CAIPI | Controlled aliasing in parallel imaging |
| CNN | Convolution Neural network |
| PE | Phase encoding |
| FE | Frequency encoding |
| PSF | Point spread function |
| SR | Super resolution |
| FOV | Field of view |
| FC | Fully-connected |
| SSIM | Structural consistency index |
| RMSE | Root Mean Squared Error |
| DC | Distortion correction |
| NEX | Number of excitations |
| $R_{PE}$ | Acceleration factor along PE direction |
| $R_{psf}$ | Acceleration factor along PSF direction |

